# BtuJ1, a novel surface-exposed B_12_-binding protein in *Bacteroidetes*, functions as an extracellular vitamin reservoir that enhances fitness

**DOI:** 10.1101/2025.07.01.662609

**Authors:** Javier Abellon-Ruiz, Raul Pacheco-Gomez, Jessica Watts, Adam Hart, Robert H. Hirt, Arnaud Baslé, Bert van den Berg

## Abstract

The acquisition of vitamin B_12_ and related cobamides is a key determinant for the fitness of *Bacteroidetes* in the gut. Depending on the species, this uptake process relies on one to four transport systems centered on conserved core outer membrane (OM) complexes composed of the TonB-dependent transporter BtuB and the surface-exposed lipoprotein BtuG. Additionally, the surface-exposed lipoprotein BtuH, although not tightly associated with the BtuBG complex, contributes to cobamide uptake and provides a fitness advantage. Here, we report the functional and structural characterization of BtuJ1 from *Bacteroides thetaiotaomicron (B. theta)*, a novel B_12_-binding lipoprotein. Under limiting B_12_ conditions, BtuJ1 is the most abundant component among the three B_12_-transport systems encoded by *B. theta*. BtuJ1 is surface exposed and binds vitamin B_12_ and cobinamide (an intermediate in B_12_ biosynthesis) with low nM affinity, conferring a fitness advantage in B_12_-limited environments. *In vitro* B_12_ transfer experiments suggest a role for BtuJ1 as an extracellular reservoir for B_12_, expanding the functionalities of the diverse group of accessory OM proteins employed by *Bacteroides* to scavenge this essential cofactor in the competitive environment of the human gut.

**IMPORTANCE:** Understanding how key molecules support bacterial colonization of the human GI tract is essential to rationalize the structure of the complex microbial community inhabiting the distal gut. The Bacteroidetes are one of the dominant phyla in this environment. Given that most Bacteroidetes cannot make vitamin B_12_ but depend on it for growth, the fitness of many species likely depends on the acquisition of vitamin B_12_. Unlike the classical model bacterium *Escherichia coli*, which encodes a single OM B_12_ transporter, the genomes of Bacteroidetes often encode multiple uptake systems comprising a heterogeneous repertoire of surface-exposed lipoproteins. These proteins may assist OM assemblies for vitamin B_12_ transport that provide fitness advantages *in vivo*.

## INTRODUCTION

The human gut microbiome is a highly complex community of microorganisms that has been linked to many aspects of health (1, 2). The vast majority of these commensal microbes inhabit the distal gut, where Bacteroidetes and Firmicutes are the dominant phyla (3). Belonging to the Bacteroidetes, the genus Bacteroides is the most abundant genus in the gut (3). With bacterial densities of 10^12^/g of luminal content(4), competition for resources is likely severe. Therefore, proficiency in the uptake of key small molecules likely is an important factor in determining competitiveness of a microorganism within the gut (5, 6). Despite this, how such small molecules are taken up in the gut is poorly understood, particularly for non-Proteobacteria, hindering the understanding of how these molecules shape the gut microbiome.

Vitamin B_12_ (cobalamin) is an organometallic cofactor that belongs to the family of cobamides and the most complex vitamin (7). It is involved in a wide variety of metabolic processes and for many organisms it is essential for the final enzymatic reaction of the L-methionine biosynthesis pathway (8, 9). Despite its key role in eukaryotic and prokaryotic cells, only a small group of microorganisms is able to produce cobalamin. Its production is expensive and requires 30 enzymatic reactions (10). Hence, microorganisms have developed dedicated systems to take up this precious resource. Bacteroidetes lack the ability to synthesise cobamides (11) and frequently code for multiple uptake systems for B_12_ and related cobamides (12), and which determine bacterial fitness in a community-and diet-dependent manner (5, 6, 12).

Escherichia coli (*E. coli*), the classical model for bacterial vitamin B_12_ uptake, possess a OM TonB-dependent transporter, BtuB, that allows the translocation of B_12_ from the extracellular space to the periplasm (13, 14), where the vitamin is bound by BtuF and transported to the cytoplasm via the ABC-transporter BtuCD (15). In contrast, gut *Bacteroidetes* often have several transporters, located in different *btu* loci (12). These loci contain the universally conserved core elements BtuB and BtuG (12, 16). These proteins form an OM protein complex with BtuG bound to the extracellular region of BtuB, acting as a mobile “lid” to assist with B_12_ uptake (17). Additionally, these loci include a heterogeneous array of genes that code for BtuCDF and for proteins with no homologues in E. coli (12). The reference species *Bacteroides thetaiotaomicron* (*B. theta*) has three B_12_ transport loci (Fig. 1A), showcasing many of these genes absent in *E. coli*. Locus 2, the most relevant *in vivo*, codes for BtuH2 (18), a surface-exposed lipoprotein that binds B_12_ and confers a fitness advantage for *B. theta* at low concentration of B_12_, but does not tightly associate with the BtuBG complex(17). B_12_ represses the expression of these loci via riboswitches but surprisingly, the abundance of the product *BT1491*, one gene from locus 1, remains high independently of B_12_ concentration (17, 19). In addition, BT1491 is the most abundant protein across the three loci (17). These findings piqued our interest and we decided to characterise this protein further.

**FIG 1.**
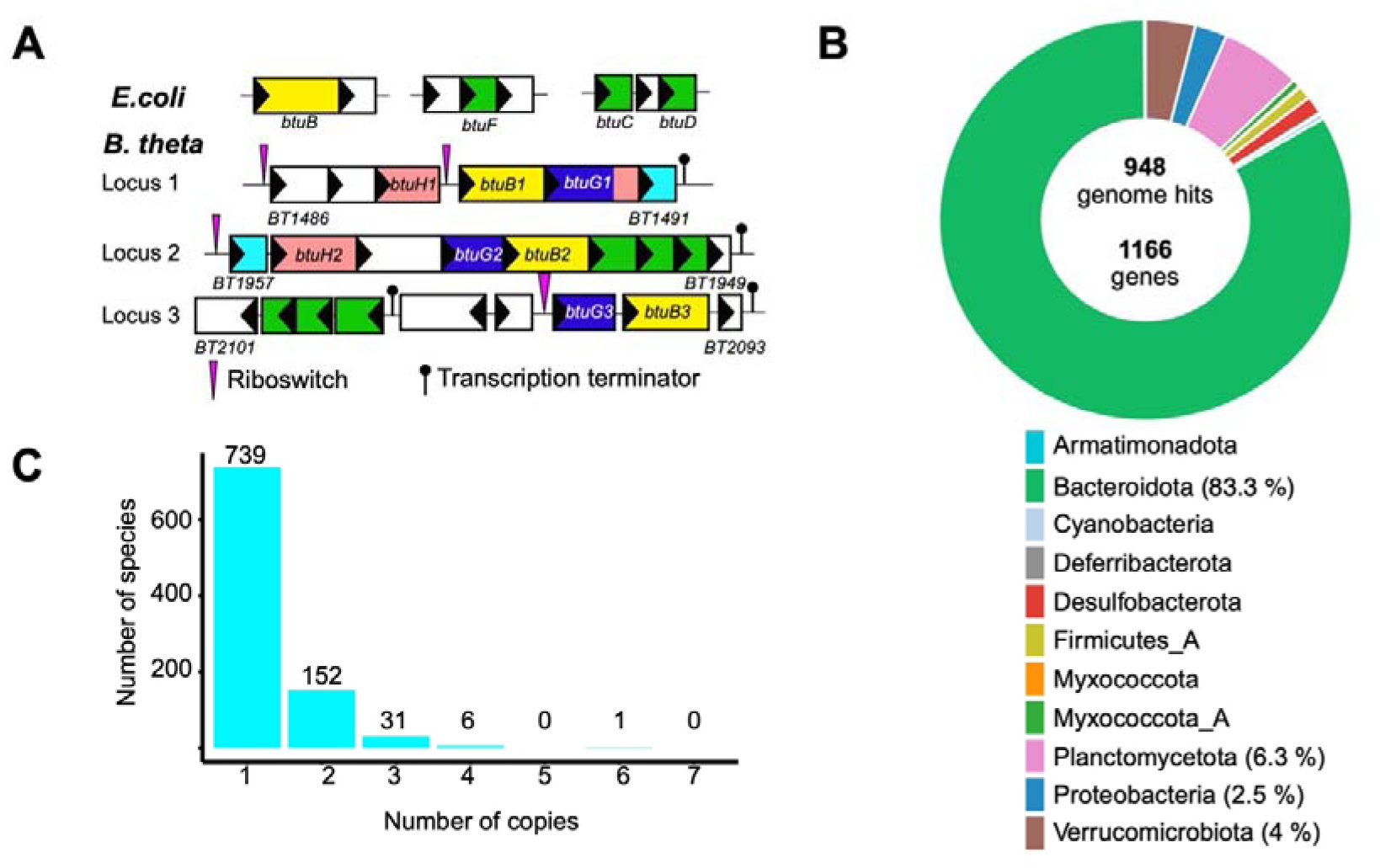
Genomic organisation of the three *btu* loci of *B. theta* and distribution and copy number of BtuJ (Pfam PF14717) orthologues across bacteria taxa. (A) Genetic organisation of the B_12_ transport system in *E. coli* (btuBFCD) and the three homologous loci in *B. theta*, showing the locations of BtuB (yellow), BtuG (blue), BtuH (pink) and BtuJ (cyan) proteins. Notice that BtuG1 has a BtuH domain fused to BtuG. The pink triangles represent B12-dependent riboswitches and the black lollipops transcription terminators. The inner membrane ABC transporters are in green. (B) Donut chart representing the number of PF14717 hits (n = 948) at the phylum level. The total number of genes detected was 1166. Each segment corresponds to the phylum shown in the legend. Genome percentages over 2.5 % are in parentheses. (C) Bar chart showing the distribution of PF14717 copy numbers per species. Each bar represents a copy number category, with the number of species possessing that copy indicated above each bar (20, 21).

Here, we show that BT1491, which due to its location in locus 1 we have named BtuJ1, adopts a jellyroll fold that binds vitamin B_12_ and cobinamide (an intermediate in B_12_ biosynthesis) tightly. *In vitro*, BtuJ1 enhances the fitness of *B. theta* under limiting B_12_ conditions and transfers B_12_ to other surface-exposed components encoded in locus 2, suggesting functional crosstalk between distinct *btu* loci and highlighting the complexity of vitamin B_12_ acquisition in human gut *Bacteroides*.

## RESULTS

### BtuJ1 homologues are widespread across Bacteroidetes

BT1491 (BtuJ1) contains a domain of unknown function, DUF4465, which is predominantly found in Bacteroidetes from the human gut microbiota, but is also present in some environmental and aquatic bacteria, including members of the phylum Planctomycetota (PF14717). A paralogue of BtuJ1 in B. *thetaiotaomicron*, BT1957 (hereafter referred to as BtuJ2), is located within locus 2 (Fig. 1A). Both BtuJ1 and BtuJ2 harbour the DUF4465 domain and share 27% sequence identity and 48% similarity, indicating relatively low sequence conservation and suggesting that these are distant homologues. Nevertheless, their moderate similarity, shared domain architecture, and genomic localization within cobamide uptake loci support a conserved function. To further investigate the distribution and copy number of BtuJ1-like proteins across microbial taxa, we conducted a phylogenetic analysis using AnnoTree (v1.2) (20). This web-based platform incorporates over 30,000 bacterial and 1600 archaeal genomes, organised according to the Genome Taxonomy Database (GTDB) (21), providing a standardised and curated taxonomic framework. The analysis identified 1,166 genes assigned to the PF14717 protein family, distributed across 948 bacterial genomes (Fig. 1B and Table S1). No PF14717 homologs were detected in archaeal genomes. The most represented phylum is Bacteroidota, accounting for ~ 83 % of all genomes harbouring PF14717 (Fig. 1B). At the family level, Bacteroidaceae comprise ~ 20 % of all genomes. Analysis of gene copy number per species revealed that 63 % of genomes carried a single copy, while 26.1 % contained two copies, suggesting that gene duplication is relatively common within this protein family (Fig. 1C).

### BtuJ1 is a surface exposed lipoprotein that confers a fitness benefit *in vitro* under limiting conditions of B_12_

As *B. theta* has three cobamide transport systems with two BtuJ paralogs, we first generated a simplified background deleting locus 2 and genes *Bt2093* to *Bt2095* from locus 3. This strain, that we call parental, contains only one BtuJ homolog (BtuJ1), one BtuBG transporter (from locus 1) and one ABC transporter (BtuFCD from locus 3; note that locus 1 lacks its own ABC transporter) (Fig. 1A). To study the fitness effect of BtuJ1 we cultured the parental strain, Δ*btuJ1*, and a complemented strain in minimal media with replete (40 nM) and limiting (0.4 nM) B_12_. The three strains show identical growth in the replete media while in B_12_-limiting media the absence of BtuJ1 abolishes growth. The complemented strain restores growth, albeit not at the level of the parent strain (Fig 2A). It is worth mentioning that the reintroduction of *btuJ1* is via the pNUB2 plasmid (22) which is integrated in a tRNA site and not in the original locus 1, providing a potential explanation for the incomplete complementation. To further investigate the importance of BtuJ1 for *B. theta* we carried out competition assays between the parental and Δ*btuJ1* strains in minimal media with replete and limiting B_12_ conditions. As shown in Fig. 2B, the parental strain rapidly out-competes the Δ*btuJ1* strain during the first 24 hours of growth.

**FIG 2.**
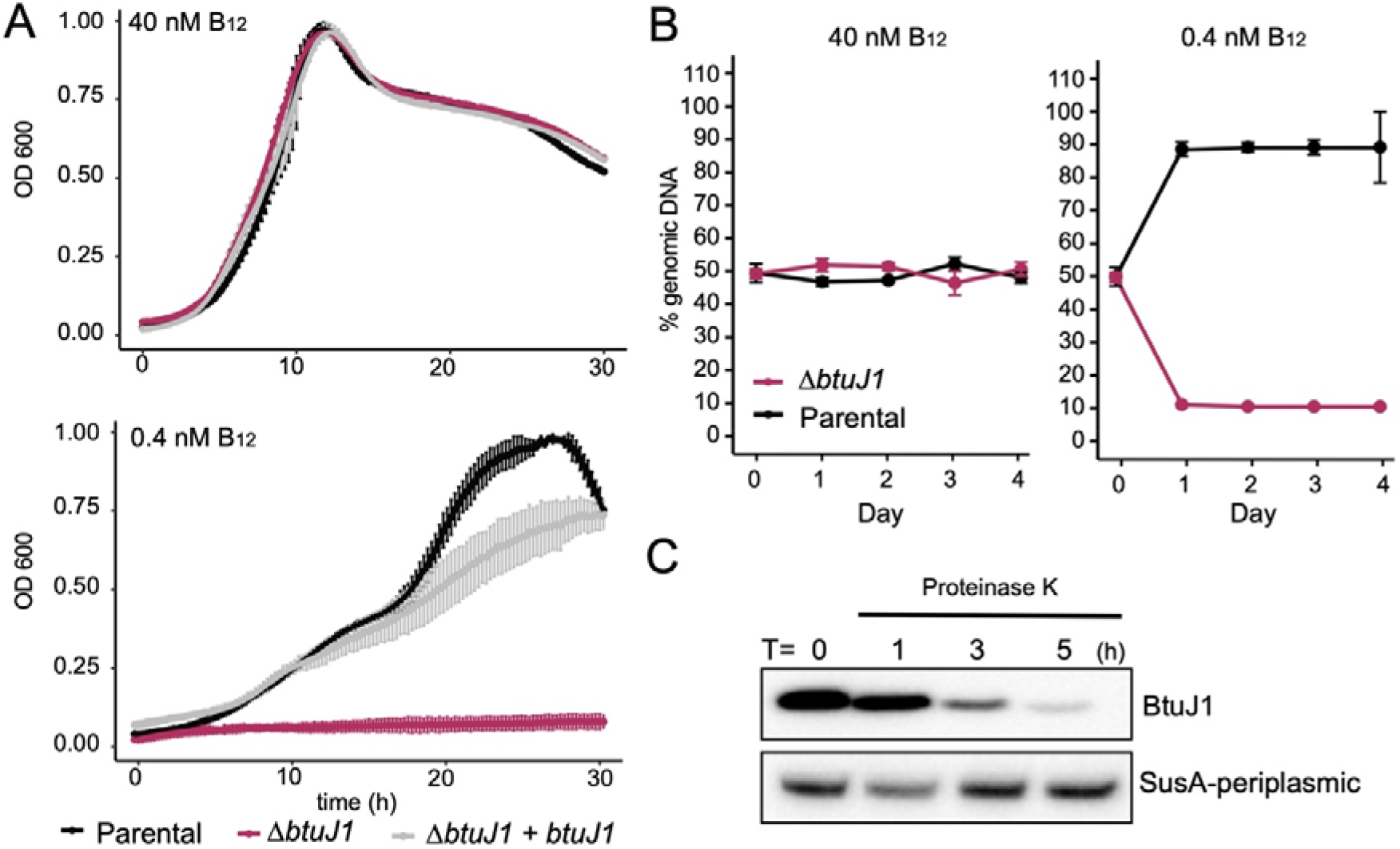
BtuJ1 is a surface-exposed lipoprotein that contributes to *B. theta* fitness in B_12_-limiting conditions *in vitro*. (A) Growth curves of several strains in minimal media with 40 or 0.4 nM vitamin B_12_. BtuJ1 is required for growth in B_12_-limiting conditions. (B) A competition assay under limiting B_12_ conditions shows that BtuJ1 confers a marked fitness advantage to *B. theta*. (C) Western blot showing that BtuJ1 is a surface-exposed lipoprotein. All panels are representative of three independent experiments. Standard deviation bars are represented in A and B.

BtuJ1 is a 256-amino-acid protein containing a predicted Sec/SPII signal sequence, which directs export via the Sec translocon and is cleaved by Signal Peptidase II (23) at a site between residues 21 and 22. This processing yields a mature 235-amino-acid peptide with an estimated molecular weight of 26.6 kDa. (All residue numbering in this text refers to the full-length peptide, with the initiating methionine as amino acid number 1). It also features the characteristic *Bacteroides* lipoprotein export signal (LES)(24) which is enriched in negatively charged amino acids in a ~6-residue window downstream of the lipidated cysteine (LES sequence of BtuJI in *B. theta* is CSDDDE, similar to the LES sequence predicted for surface-exposed lipoproteins in the closely related species *B. fragilis* (CSDDDD)) (24). To ascertain if BtuJ1 is a surface exposed lipoprotein, as the amino acid sequence suggests, we generated a strain with a C-terminal hexa-histidine tag in BtuJ1, incubated intact cells grown in B_12_-limiting minimal media with proteinase K for several hours, and probed by Western blot using anti-His antibody. This was done in parallel with a strain expressing His-tagged SusA (BT3704, a periplasmic protein) (25). The BtuJ1 signal was degraded over time while SusA was protected from degradation by Proteinase K, in agreement with BtuJ1 being a surface-exposed lipoprotein (Fig. 2C).

### BtuJ1 adopts a jellyroll fold

Crystallization of BtuJ1 in complex with cyanocobalamin (CNCbl) and the intermediate cobinamide (Cbi) (Fig. 3A) was successful, enabling structure determination using data to 1.4 Å resolution (Table S2 and Movie S1). The electron density permitted modelling from residue Met29 to the C-terminal Lys of the mature protein. BtuJ1 adopts a compact jellyroll fold, stabilized by antiparallel N-terminal and C-terminal β-sheets that interlock the structure (Fig. 3B and Fig S1A). The distal region is loop-rich, forming a negatively charged clamp-like binding site (Fig. S2A). High-quality electron density enabled unambiguous modelling of both ligands (Fig SC and D). Analysis with PDBe PISA (26) identified 15 hydrogen bonds involving 12 protein residues and the amide functional groups of substituents at positions C2, C3, C7, C8, and C18 of the corrin ring, indicating a conserved and specific binding mode (Fig 3C and Fig S1B). B_12_ is positioned within the binding cleft such that the lower ligand projects outward, making no significant interactions with BtuJ1— apart from a hydrophobic contact with Tyr97—while the upper ligand faces inward towards a small cavity (Fig. S2B). BtuJ1 also binds adenosylcobalamin, which has a bulkier upper ligand compared to other cobalamins (Fig. S3A). Attempts to obtain a crystal structure of BtuJ1 bound to adenosylcobalamin (AdoCbl), were unsuccessful. To gain structural insight into how AdoCbl fits within the confined binding cavity, we used Chai-1 (27) to model the BtuJ1–AdoCbl complex. The resulting structure revealed that AdoCbl occupies the same binding pocket as cyanocobalamin (CNCbl), but with a notable difference: the corrin ring of AdoCbl is rotated by approximately 100 degrees, although it remains in a similar plane. Additionally, binding of AdoCbl induces conformational rearrangements in seven BtuJ1 residues. In particular, the side chains of Trp78, Tyr57, and Glu55 undergo substantial displacement, effectively enlarging the upper ligand cavity to accommodate the bulky adenosyl group (Fig. S3B). A ConSurf analysis performed using 250 homologues revealed moderate overall residue conservation across BtuJ1. Importantly, residues involved in hydrogen bonding with vitamin B12 were among the most conserved, confirming functional importance (Fig. S4). While BtuJ1 and its paralogue BtuJ2 in *B. theta* share only 27% sequence identity and 48% similarity, structural superposition of AlphaFold-predicted models revealed a similar fold, with an RMSD of 1.65 Å across 228 aligned Cα atoms as analysed in COOT (28). This structural conservation, despite limited sequence similarity, supports the notion of a preserved core function between these distant homologues.

**FIG 3.**
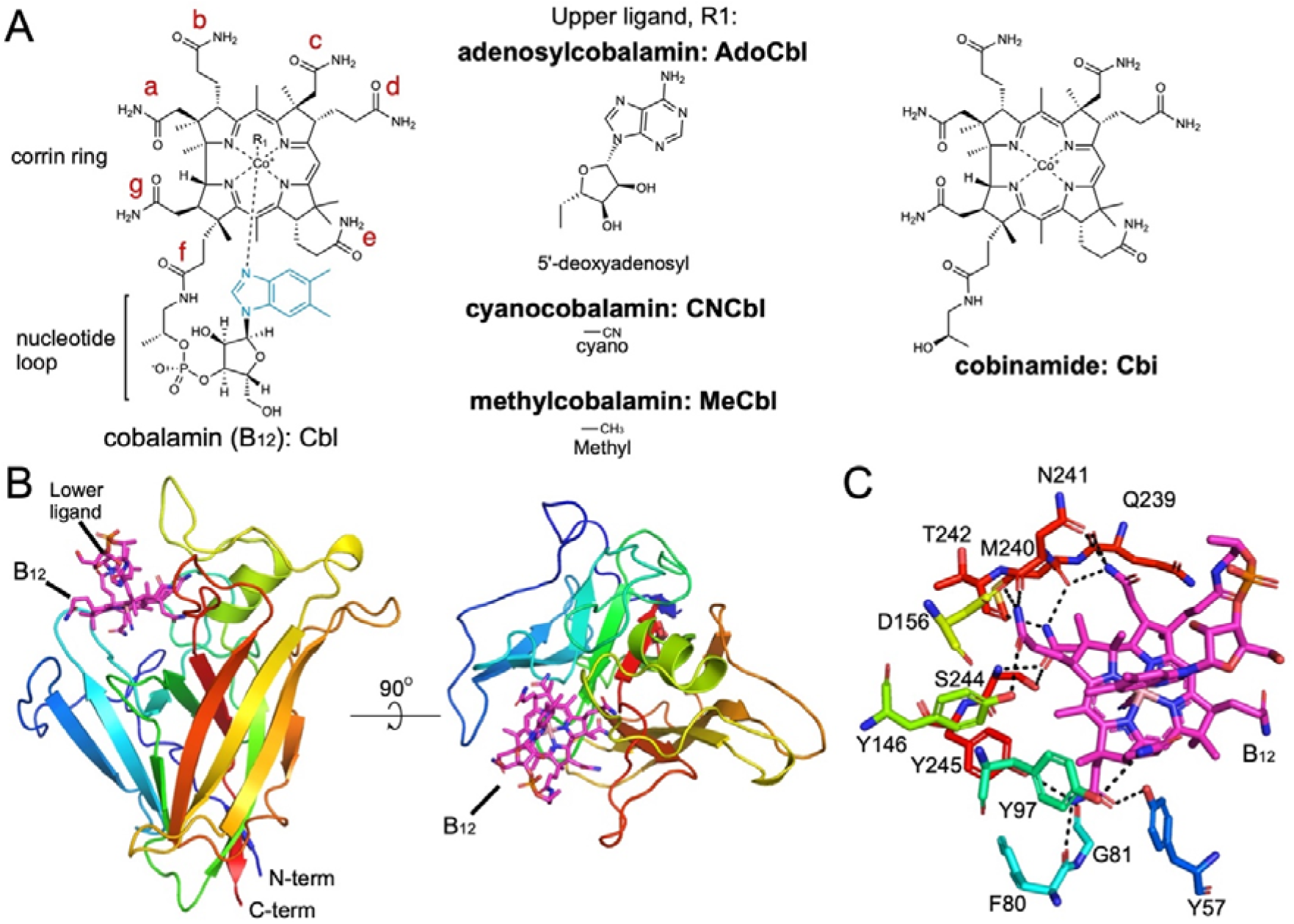
BtuJ1 adopts a jelly roll fold with a clamp-like binding pocket for different corrinoids. (A) Diagram showing the corrinoids used for crystallography (left and right). Lower ligand in blue, side chains of the corrin ring labelled with letters in red. In the central panel, the biologically active upper ligands—adenosylcobalamin (AdoCbl) and methylcobalamin (MeCbl)—are shown alongside cyanocobalamin (CNCbl), the synthetic and non-physiological form. (B) Cartoon representation of BtuJ1 (in rainbow colour, N terminus in blue) bound to CNCbl (magenta). Note that the lower ligand is pointing outwards. (C) Close-up of the residues forming hydrogen bonds (black dashed lines) with CNCbl.

### BtuJ1 binds B_12_ with nanomolar affinity

To measure the binding affinity Isothermal titration calorimetry (ITC) was used initially to assess the binding affinity of BtuJ1 for cyanocobalamin. In all experiments, the transitions in the enthalpy curves were extremely rapid, hampering fitting and indicating affinities that are at or close to the limit of ITC (low nM) (Fig. S5A). To more precisely determine the dissociation constant (K_D_) as well as k_on_ and k_off_ values, we conducted grating-coupled interferometry (GCI) using cyanocobalamin (CNCbl) and cobinamide (Cbi) as ligands at two concentrations (100 nM and 250 nM). Both ligands yielded similar dissociation constants, with CNCbl binding slightly more tightly than Cbi (3.6 nM vs. 11.4 nM at 100 nM), based on a 1:1 binding model and assuming a single site in BtuJ1. A mass transport-corrected (MT) model returned comparable K_D_ values, indicating minimal mass transport limitations (Fig. S5B and S5C). These results confirm a high-affinity interaction in the low nanomolar range, consistent with ITC observations and previous work using a B_12_-affinity-based probe to capture cobamide binding proteins in *B. theta* (18)

### BtuJ1 transfers B12 to other surface-exposed lipoproteins of locus 2

A striking feature of many gut bacteroidetes is the presence of multiple cobamide uptake systems, each encoded in separate genomic loci (12). In addition to the TonB-dependent transporter BtuB, these loci contain a diverse array of proteins involved in cobamide acquisition (16, 18). In *B. theta*, locus 2 appears to be the most functionally relevant *in vivo* and includes several surface-exposed lipoproteins that facilitate cobamide uptake—namely, BtuG2, which is universally conserved across all uptake loci (16), and BtuH2, which is found in approximately two-thirds of gut Bacteroidetes (12, 18). Our previous work demonstrated that BtuGs form stable complexes with BtuBs, but other surface lipoproteins do not strongly associate with the BtuBG complex (17). This raises the intriguing possibility that these other B_12_-binding proteins, such as BtuJ1, may function in coordination with BtuBG complexes from different loci, enabling potential cross-talk between uptake systems. To test whether BtuJ1 can transfer B_12_ to BtuG2 and BtuH2 from locus 2, we performed cobamide transfer experiments (Fig. 4A). In these *in vitro* assays, apo proteins (lacking the lipid anchor) were incubated with B_12_-loaded partner proteins, with no free B_12_ present in the buffer—ensuring that any transfer must occur directly between proteins. Protein elution profiles were monitored via absorbance at 280 nm (protein) and 362 nm (CNCbl) (Fig. 4B and Fig. S6), allowing us to determine whether B_12_ remained bound to the donor protein or was transferred to the recipient. The results show that when B_12_-bound BtuJ1 is incubated with apo-BtuG2 or apo-BtuH2, B_12_ is efficiently transferred—100% and ~ 80%, respectively. In contrast, when BtuG2 or BtuH2 are the B_12_ donors, no transfer to BtuJ1 was observed (Fig. 4C). These findings suggest that BtuJ1 can donate B_12_ to locus 2-encoded proteins, at least *in vitro*.

**FIG 4.**
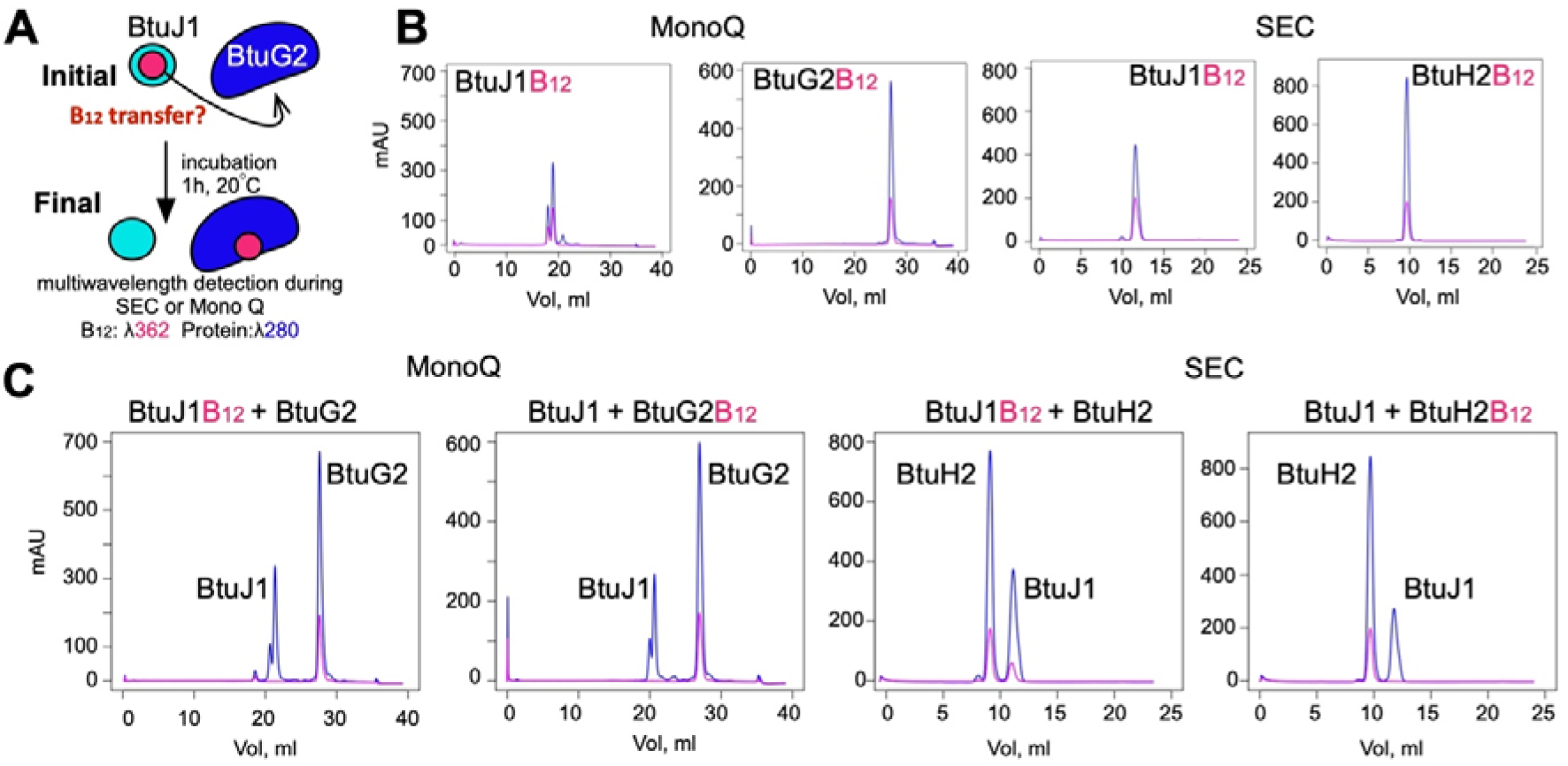
CNCbl (vitamin B_12_) transfer assays among BtuJ1 and locus 2 proteins BtuH2, and BtuG2. (A) Schematic of the transfer assay: protein elution is monitored by absorbance at 280 nm (protein) and 362 nm (CNCbl). (B) Control chromatograms for individual CNCbl-bound proteins (5 nmol each), on Mono Q ion-exchange (left) and Superdex 75 SEC (right). Each trace is labelled with the corresponding protein. (C) Chromatograms from co-incubation of CNCbl-loaded protein with apo protein (indicated above each panel), analysed by Mono Q (left) and SEC (right). Traces are representative of two independent experiments.

## DISCUSSION

Successful colonization of the human distal gut by bacteria depends on a combination of microbial traits and host-microbe interactions that promote persistence, competition, and adaptation in a densely populated and highly competitive environment. One key determinant is metabolic versatility, particularly in nutrient acquisition. *Bacteroides* species, for example, can access a wide array of carbon sources due to an extensive repertoire of polysaccharide utilization loci (PULs), enabling the degradation of both dietary and host-derived glycans (29, 30). In such a competitive microbial ecosystem, efficient scavenging of essential metabolites becomes crucial. This gives rise to a “secondary economy” of shared, growth-limiting compounds, such as vitamins like B_12_, that shape microbial fitness and colonization success (5, 6, 31).

Bacteroidetes are cobamide consumers, unable to synthesize B_12_ de novo, yet they depend on these cofactors for survival. Their ability to outcompete others for B_12_ likely plays a pivotal role in their persistence in the gut. But how do they gain this competitive edge? Most Bacteroidetes encode multiple B_12_ transport systems, three in *B. theta*, each associated with surface-exposed lipoproteins that likely enhance uptake efficiency. This strategy resembles the utilisome, the glycan uptake machinery, where surface lipoproteins form stable complexes with TonB-dependent transporters (32). However, B_12_ and glycan uptake differ in one important respect: only BtuG forms a stable complex with BtuB in B_12_ transport (17). In contrast, other B_12_-binding lipoproteins, like BtuH2, do not stably associate with BtuBG complexes (17). This difference may reflect the nature of the substrates. Glycans are large molecules requiring coordinated binding and enzymatic processing, favouring multi-component complexes. In contrast, vitamin B_12_ is small and scarce, making efficient capture from a competitive environment the central challenge, rather than its processing.

In this work, we demonstrate that BtuJ1, an additional surface-exposed lipoprotein that binds B_12_, confers a competitive advantage under low B_12_ concentrations. Alongside BtuGs and BtuHs, BtuJ1 assists in B_12_ uptake, highlighting the high value of this cofactor to *Bacteroides*. Remarkably, *B. thetaiotaomicron* encodes at least 16 dedicated proteins in the cell envelope to acquire this molecule. We cannot rule out that other genes within the three B_12_ uptake loci also contribute to B_12_ binding or assist indirectly, adding layers of complexity to an already intricate system. But perhaps ligand size and scarcity are not the only factors determining whether lipoproteins are integrated into a complex or act independently. It is tempting to propose that “freelance” lipoproteins, such as BtuJ1, may function across loci and not just with their native BtuBG partners. In this model, a bacterium expressing BtuJ1 could cooperate with the BtuB2G2 system to enhance B_12_ acquisition, providing a fitness advantage and a possible evolutionary rationale for the retention of multiple transporters within the same species. Our *in vitro* data support this hypothesis: BtuJ1 can transfer B_12_ to BtuH2 and BtuG2, spanning affinity ranges from low nanomolar to femtomolar/sub-picomolar, respectively. This suggests a mechanism resembling a “bucket brigade”, where mobile, surface-exposed lipoproteins hand off B_12_ to the nearest available transporter, rather than seeking a specific complex. Given the limited mobility of outer membrane proteins in Gram-negative bacteria such as *E. coli*, transferring B_12_ to the closest partner may be the most effective strategy.

In our previous work (17), we showed that BtuJ1 exhibits a unique feature that distinguishes it from other proteins in the B_12_ uptake system, including its paralogue BtuJ2. Typically, B_12_ uptake loci are negatively regulated by intracellular B_12_ levels: high concentrations repress gene expression via riboswitches, leading to a gradual decrease in the abundance of associated proteins in the outer membrane (OM). However, BtuJ1 defies this pattern—its levels in the OM remain high regardless of B_12_ concentration. What might be the biological significance of this? One possibility is that BtuJ1 serves a dual function: first, as a scavenger, sequestering B_12_ from the environment to limit access by competing species; and second, as a reservoir, retaining B_12_ at the surface for future use by B. *thetaiotaomicron*. While intracellular B_12_ still suppresses transcription of the uptake machinery, BtuJ1 continues to capture external B_12_. Then, when internal levels drop and transcription is reactivated, BtuJ1 may act as the first link in a “bucket brigade”, immediately transferring B_12_ to newly synthesized uptake components as they are inserted into the OM (Fig. 5).

**FIG 5.**
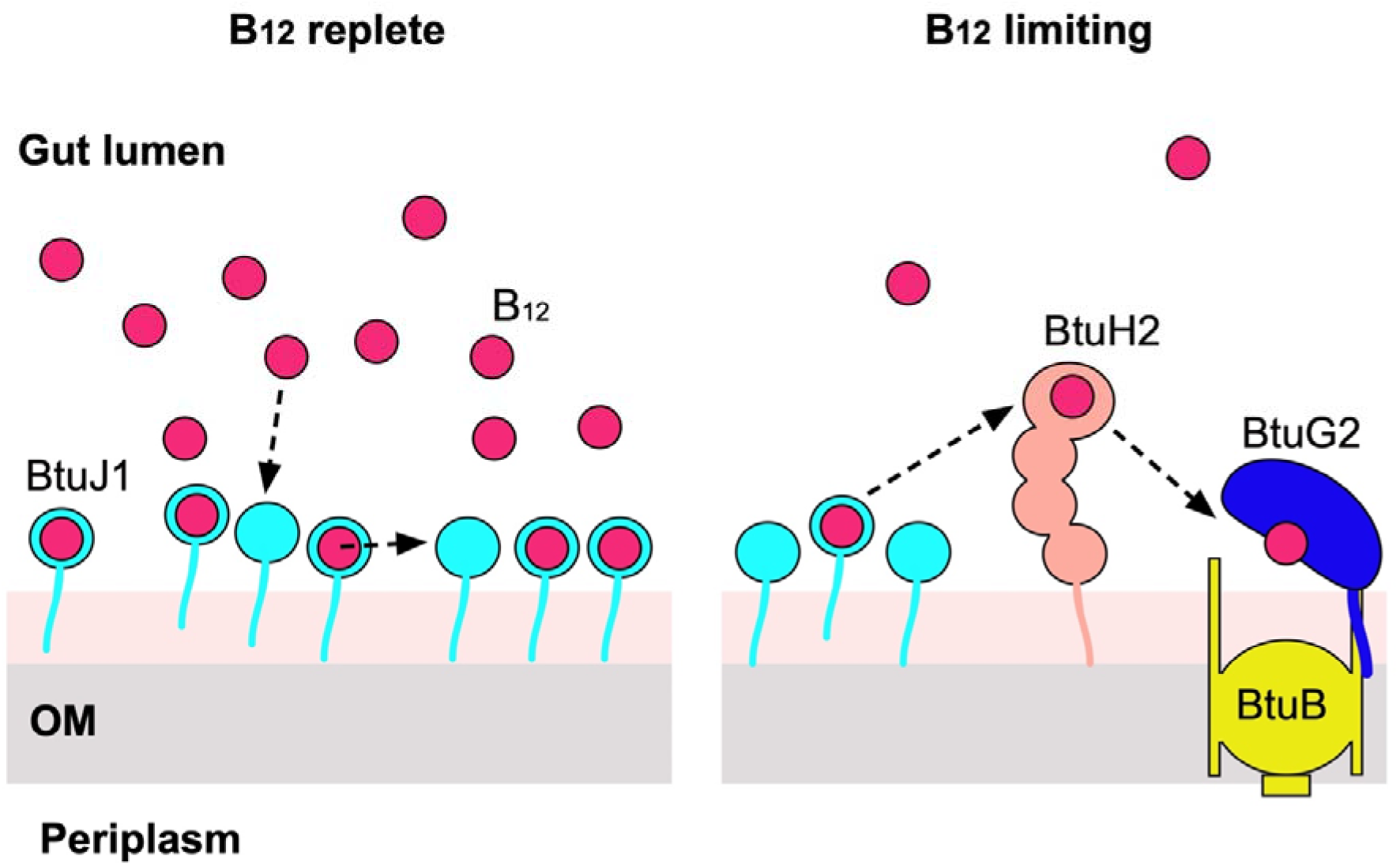
Schematic model for lipoprotein-mediated B_12_ acquisition by *B. theta*. In a B_12_-replete environment, BtuJ1 is the most abundant B_12_-binding protein among the three uptake loci encoded by *Bacteroides thetaiotaomicron*. Localized in the outer membrane (OM), BtuJ1 scavenges and stores B_12_ from the gut lumen, effectively functioning as a B_12_ “pantry.” When intracellular B_12_ levels decline, the expression of other uptake components is induced, and they are inserted into the OM. In this context, BtuJ1 serves as a readily available reservoir of B_12_, facilitating its transfer to BtuBG complexes for transport into the cytoplasm.

Together, our findings highlight the complexity and specialization of cobamide acquisition systems in *Bacteroidetes*. The presence of multiple uptake loci, the differential behaviour of surface-exposed lipoproteins like BtuJ1, and the apparent capacity for functional cooperation across loci point to a modular and adaptive strategy for micronutrient scavenging in a competitive environment. These features likely contribute to the ecological success of *Bacteroidetes* in the human distal gut, where access to essential but scarce resources such as vitamin B_12_ can define fitness. Understanding how such “bucket brigade” mechanisms function *in vivo* offers exciting opportunities for future research into microbial nutrient economies and interbacterial interactions in the gut microbiome.

## MATERIAL AND METHODS

### Culture conditions

*B. thetaiotaomicron* VPI-5482 was routinely grown on brain-heart infusion (BHI, OXOID) supplemented with hemin (1 μg/ml, Sigma-Aldrich). Agar (Fisher bioreagents), erythromycin (25 μg/ml, Duchefa Biochemies), gentamicin (200 μg/ml, Formedium) and 5-fluoro-21-deoxyuridine (FUdR, 200 μg/ml, Sigma-Aldrich) were added when required. Minimal medium (MM)(22) contained 7.5 mM NH_4_SO_4_ (Thermo Scientific), 9.5 mM Na_2_CO_3_ (Melford), 4 mM L-Cysteine (Melford), 100 mM KH_2_PO_4_ (pH 7.2) (Fisher Chemical), 1 μg/ml menadione (Sigma-Aldrich), 1 μg/ml hemin and a mixture of mineral salts (15 mM NaCl (Duchefa Biochemies), 0.18 mM CaCl_2_•2H_2_0 (Fisher Chemical), 0.1 mM MgCl_2_•6H_2_O(Fisher Chemical), 0.5 mM MnCl_2_•4H_2_O (Acros Organics) and 0.04 mMCoCl_2_•6H_2_O (Sigma-Aldrich)). Vitamin B_12_ (in the form of CNCbl, Sigma-Aldrich) and L-methionine (Formedium) were added as required and described in the text. 0.5% fructose (Thermo scientific) was added as carbon source. Cells were grown in a Whitley A35 anaerobic workstation with a gas mixture of 80% N_2_, 10% CO_2_ and 10% H_2_. Table S3 contains a list with the strains, plasmids and primers used in this work.

### Cloning, expression and protein purification

The coding region for the mature part of BT1491 (btuJ1) excluding the anchor methionine was cloned into pET28 using NcoI and XhoI restriction sites. The protein was expressed in BL21(DE3) in LB broth (Lennox L. Broth, Melford) 2.5 h at 37 °C with the addition of 1 mM IPTG (Melford) at OD ~ 0.6 to induce protein expression. Cells were collected by centrifugation, resuspended in TBS (10 mM Tris (Sigma-Aldrich)), 300 mM NaCl; pH 8) lysed at a pressure of 20–23 kpsi using a cell disrupter (Constant Systems 0.75 kW), and purified by nickel affinity chromatography (Briefly, the lysed pellet was centrifuged for 30 minutes at 4 °C at 19,000 rpm in a JA25.50 rotor, the supernatant loaded onto a 1 ml Ni-charged chelating sepharose fast flow (GE Healthcare), washed with 20 column volumes (CVs) TBS plus 30 mM Imidazole and eluted increasing the imidazole to 250 mM in 2.5 CVs) in TBS buffer. Protein concentrations were measured using the BCA assay, and when needed, cyanocobalamin (CNCbl), dicyanocobinamide (Cbi) or adenosylcobalamin (AdoCbl, all from Sigma-Aldrich) were added to a molar ratio 1:2 protein:corrinoid. The samples were incubated at 4 °C for 30 minutes and subjected to SEC using a HiLoad 26/600 Superdex200 column (GE healthcare) in 10 mM Hepes, 100 mm NaCl, pH 7.5.

### Bioinformatic analysis

Genomic occurrence data for Pfam protein family PF14717 were retrieved from AnnoTree (V 1.2) (20). Taxonomic assignments were matched using the Genome Taxonomy Database (GTDB), release R95 (21) to ensure consistency in phylogenetic classification. Hit counts were computed at the phylum and species levels and plots showing the relative abundance of PF14717 across phyla (donut chart) and number of hits per species (bar chart) were generated using the R package (Version 4.2.3) Ggplot2 (Version 3.4.2)(33). The model of BtuJ1-AdoCbl was generated using the protein sequence and adenosylcobalamin as the ligand, using the Chai Discovery webtool (27).

### Crystallisation and X-ray crystal structure determination

Sitting drop vapour diffusion crystallization were set up with a Mosquito Crystallization robot (TTP Labtech) using the commercial screens Index (Hampton research), Structure, JCSG+, Morpheus and PACT (Molecular Dimensions). Screens were performed at 20 °C. Crystals obtained in Index D4 (0.1 M citric acid pH 3.5, 25% PEG 3350) from BtuJ1-B_12_ and BtuJ1-Cobinamide were cryoprotected with 20 % PEG 400 and flash-frozen in liquid nitrogen. Diffraction data were collected at 100 K at the Diamond Light Source (supplementary Table 2). Both datasets were integrated with XDS(34), the space group determined with Pointless (35), and data were truncated (CC½ ≥ 0.5) and scaled with Aimless. For BtuJ1-B_12_ the phase problem was solved via Co-Sad autosolve(36, 37) and the resulting model was used for molecular replacement in phaser(38) for the BtuJ1-Cbi dataset. Autobuild(39) was used to generate an initial model, improved with subsequent rounds of manual building in coot(28) and phenix refine(40). B_12_ and Cbi molecules were added from the REFMAC monomer library using codes CNC and CBY respectively. MolProbity(41) was used to validate protein geometry and PyMOL was used of the visualization of the protein structures. ChimeraX was used to generate the move file (42). Residue conservation was analysed using ConSurf with default parameters, except for the number of homologous sequences, which was increased to 250(43).

### Proteinase K assay

10 ml of MM (0.4 nM of B_12_) per time point was inoculated with an overnight culture of *B. theta* Δ*tdk btuJ1*-6xHis to an OD_600_=0.04 and grown in anaerobic conditions to OD_600_=0.8. Cells were pelleted and stored at −20 °C. Samples were washed with PK buffer (50 mM Tris, pH 7.5, 50 mM NaCl and 1 mM MgCl), resuspended in the same buffer with or without 500 ug/ml proteinase K and incubated at 37 °C for 0, 1, 3 and 5 hours. Cells were washed twice with PK buffer containing 2 mM phenylmethylsulfonyl fluoride (PMSF), and proteins solubilised in BugBuster protein extraction reagent (Millipore), incubated at 25 °C for 25 minutes with vigorous shaking, spin down (14,000 rpm) in a microcentrifuge for 10 minutes and subjected to SDS-polyacylamide gel electrophoresis (PAGE). The gel was analysed by western blot using Roche anti-his6-peroxidase (as per the manufacturer’s specifications) and developed with SuerSignal West Pico Chemiluminiscent Substrate (Thermo Scientific). Chemiluminescent activity was recorded with the ChemiDoc Imaging System (Bio Rad). To generate the control strain, a 6 x his-tag was added to the C-terminus of genomic *bt3704* (*susA*) using pEchange-tdk(44).

### Growth curves

To generate the parental strain, deletions of *btu* locus 2 and genes *bt2093-94-95* (from locus 3) were sequentially introduced into *B. theta* Δ*tdk*(44) (*bt2275*) using the plasmid pEchange-tdk(44). In this background, *btuJ1* was subsequently deleted using the same strategy. For complementation, *btuJ1* was placed under the control of its native operon promoter and integrated in the genome of the triple-deletion strain (Δ*locus2* Δ*bt2093-95* Δ*btuJ1*) using the pNBU2(22) vector. Integration at either the *att1* or *att2* site was confirmed by PCR screening. Promoter selection was guided by the RiboD database (45) to confirm inclusion of the complete riboswitch element. Overnight cultures of the desired strains were grown overnight under anaerobic conditions in BHI supplemented with hemin (1 μg/ml), and subcultured for 4 h next morning in fresh media. Cells were washed two times with minimal media without B_12_, the OD600 was adjusted to 0.03 and 100 μl of cells were dispensed into wells of a 96-well plate. Negative control wells (without B_12_ and L-methionine), positive control wells (with 200 mM L-methionine) and cultures with permissive and restrictive conditions for B_12_ were set up in triplicates and grown for 24 h in the anaerobic cabinet. OD measurements were done using a Biotek Epoch microplate reader. Data was collected using Biotek Gen5 (2.09 Agilent) software and analysed and plotted with the R package (Version 4.2.3) Ggplot2 (Version 3.4.2)(33).

### Competition assays

Overnight cultures of the desired *B. theta* strains carrying unique oligonucleotide barcode sequences(22) (integrated at the *att1* site via the pNBU2 (22) vector) were grown overnight under anaerobic conditions in BHI supplemented with hemin (1 μg/ml). The next morning, they were subcultured in MM with L-methionine until an OD600 of 0.5 was reached. Cells were washed once with minimal media without B_12_, the OD600 was adjusted to 0.25 in MM and both competing strains combined to a total final OD600 of 0.5 (Day 0). A 1:10,000 dilution was prepared in 2 ml of MM plus 200 mM L-methionine or B_12_ (permissive and restrictive conditions), all in triplicates. Cultures were incubated anaerobically at 37 C and passaged every 24 hr by 1:1,000 dilution into fresh MM for four days. Total gDNA was recovered from cultures each day, including the initial day 0 (GenElute Bacterial Genomic DNA kit, SIGMA) and the relative abundance of each strain determined by qPCR as described(22) using a LightCycler 96 instrument (Roche) and Luna universal master mix (New England biolabs). Mean strain quantities were calculated using a standard curve and relative fold changes were analysed with the efficiency-corrected DeltaCq method (46).

### B_12_ transfer experiments

Five nanomoles of each protein of interest, in either apo and B12-bound forms, were co-incubated in pairs for 1 hour at 20 °C and subsequently separated using either size exclusion chromatography (Superdex 75 Increase 10/300 GL, Cytiva) or ion exchange chromatography (Mono Q Capto™ HiRes Q 5/50, Cytiva), depending on the size of the proteins being compared. Separation was performed using an ÄKTA system equipped with a multi-wavelength detector, with absorbance monitored at 280 nm for protein and 362 nm for CNCbl. Control runs consisting of individual apo or B12-loaded proteins were included to allow comparison with the protein mixtures.

### Isothermal Titration Calorimetry (ITC)

ITC was performed using a MicroCal PEAQ-ITC machine with v1.41 control software for data collection (Malvern Panalytical). Briefly, pure BtuJ1 (20 µM) in a 300 µl reaction well was injected 13 times with 21µl followed by 12 times 1 µl of B_12_ (140 µl) at 20 °C, all in triplicates. Integrated heats were fit to a one set of sites model using the Microcal PEAQ-ITC analysis software v1.41 (Malvern Panalytical) to obtain K_d_, DH and N (number of binding sites on the protein).

### Grating-Coupled Interferometry (GCI)

Binding kinetics for BtuJ1 were measured using a waveRAPID (Repeated Analyte Pulses of Increasing Duration) system on a Creoptix WAVEsystem (Malvern Panalytical). BtuJ1 was immobilized onto a fresh 4PCH-NTA microfluidics chip via His-guided amine coupling on a Ni^2+^-NTA-functionalized GCI surface, using the RAPID protocol to achieve oriented capture and covalent stabilization. The ligands cobinamide (Cbi) and cyanocobalamin (CNCbl) were tested at two concentrations (100 nM and 250 nM). We conducted two experiments, increasing the flow rate in experiment 2 to mitigate potential bispecific behaviour. All measurements were performed at 20 °C in 100 mM NaCl, 10 mM HEPES pH 7.5. Data were analysed using WAVE software with both 1:1 binding and mass transport-limited (MT) models.

## DATA AVAILABILITY

The data supporting the findings of this study are available upon the corresponding authors upon reasonable request. For X-ray structures, coordinates and structure factors have been deposited in the Protein Data Bank with accession codes 9QPM for Btuj1-CNCbl and 9QPN for Btuj1-CNCbi

## ACKNOWLEDGEMENTS

The research of B.v.d.B. is supported by a Wellcome Trust Investigator award (214222/Z/18/Z), providing salary support for J.A.R. We thank the Diamond Light Source for access to crystallography beamlines (i24, proposal mx24948 and i04, proposal mx32736), and Victor Hernandez-Rocamora and Mariusz Madej for their valuable input on the phylogenetic analysis.

## AUTHOR CONTRIBUTIONS

J.A.-R. expressed and purified proteins, determined X-ray crystal structures, performed the functional, phylogenetic, and isothermal titration calorimetry (ITC) analyses. R.P.-G. and J.W. conducted and analysed the grating-coupled interferometry (GCI) experiments. A.B. maintained the Newcastle Structural Biology Laboratory. A.H. and R.H.H. provided expert input on the phylogenetic analyses. B.v.d.B. supervised the structural studies. J.A.-R. and B.v.d.B. wrote the manuscript.

