## Supplementary material for "BtuJ1, a novel surface-exposed B_12_-binding protein in *Bacteroidetes*, functions as an extracellular vitamin reservoir that enhances fitness"

**SUPPLEMENTARY INFORMATION**

**Table S1** Summary of BtuJ1 (PF14717) homolog distribution across bacterial genomes based on AnnoTree results. This table compiles data retrieved from the AnnoTree webserver using the Pfam identifier. Dataset is divided into three sheets. 1: hit_distribution, contains a summary of the number of homologs detected per taxonomic group. 2: annotree_hits, list all individual hits identified including genomic identifiers, reflecting the raw output of the query. 3: genome_to_species: matches genome accession IDs to their corresponding species names in the Genome Taxonomy Database.

**Table S2** X-ray crystallographic data collection and refinement statistics. Values in parentheses are for the highest resolution shell.

|  | **BtuJ1-CNCbl** | **BtuJ1-Cbi** |
| --- | --- | --- |
| DLS beamline | i24 | i04 |
| Wavelength | 0.978 | 0.969 |
| Space Group | P2_1_ 2_1_ 2 | P2_1_ 2_1_ 2 |
| Cell dimensions |  |  |
| a,b,c (Å) | 52, 113, 43 | 40, 51, 115 |
| a,b,g (°) | 90, 90, 90 | 90, 90, 90 |
| Molecules in AU | 1 | 1 |
| Resolution range(Å) | 38-1.4 (1.42-1.40) | 57.7-1.4 (1.5-1.4) |
| I/ σI | 20.5 (1.5) | 17.2 (0.8) |
| Completeness (%) | 99.8 (96.8) | 100 (100) |
| Redundancy | 30.2 (12.9) | 13 (13) |
| Rpim (%) | 2.1 (49) | 2 (94) |
| CC (1/2) | 1 (0.55) | 1 (0.52) |
| Anomalous completeness | 99.2 (89.7) | 100 (100) |
| Anomalous redundancy | 14.9 (6.3) | 6.8 (6.7) |
| **Phasing** |  |  |
| SOLVE FOM | 0.29 | - |
| Sites found [expected] | 1, [1] | - |
| **Refinement** |  |  |
| Rwork/Rfree (%) | 17.8/20.1 | 18.9/19.7 |
| Reflections | 50351 | 41881 |
| No. Atoms |  |  |
| Protein | 1815 | 1815 |
| Corrinoid | 93 | 72 |
| B-factors (Å2) |  |  |
| Protein | 21.4 | 31.7 |
| Corrinoid | 17.0 | 31.5 |
| Rmsd |  |  |
| Bond lengths (Å) | 0.01 | 0.009 |
| Bond Angles (°) | 1.10 | 1.01 |
| Molprobity clashscore | 2.7 | 2.4 |
| Ramachandran plot |  |  |
| Favoured (%) | 97.8 | 97 |
| Disallowed (%) | 0 | 0 |
| PDB code | 9QPM | 9QPN |

**Table S3** Strains, plasmids and primers used in this work**.**

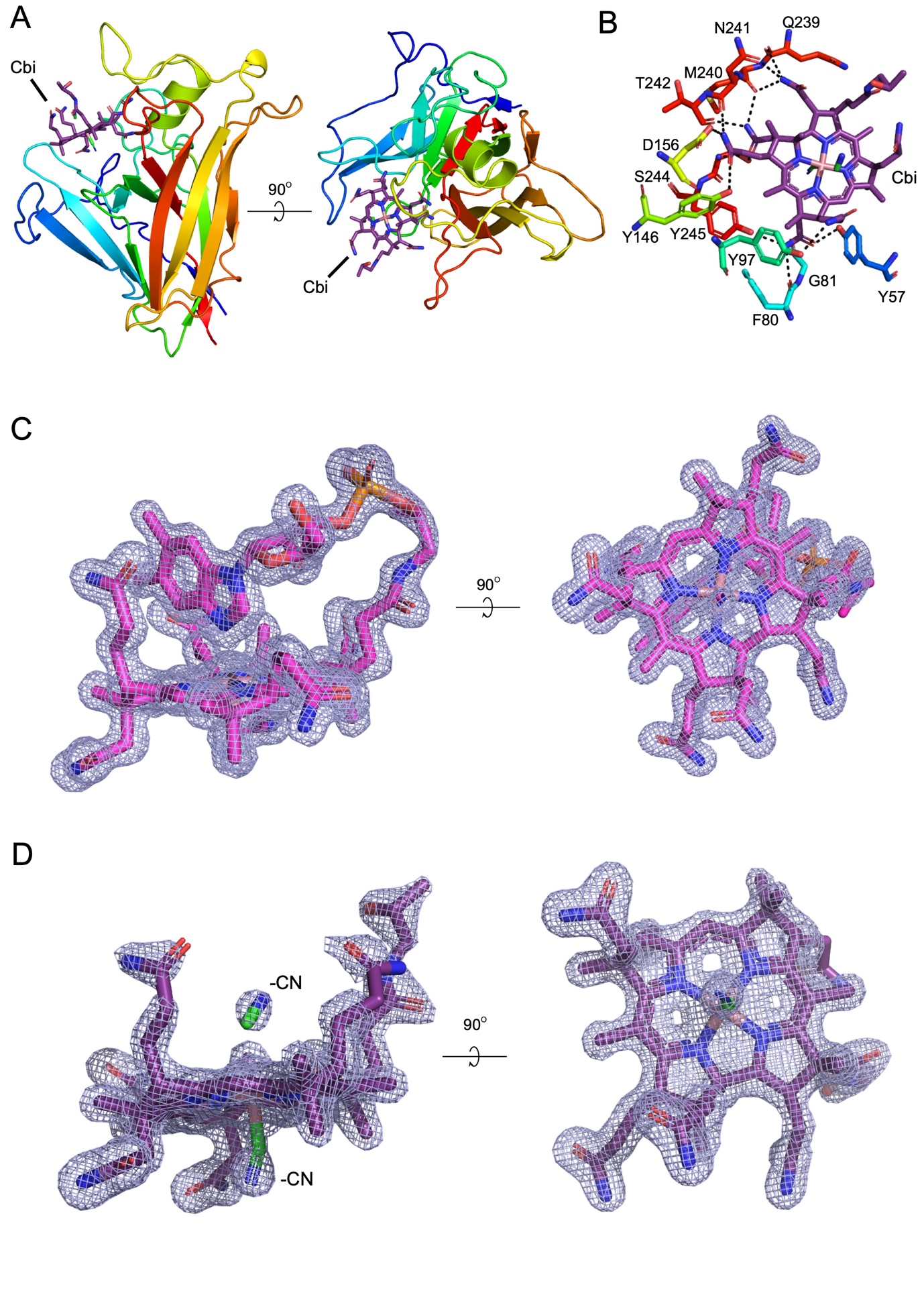

**FIG S1** Structural analysis of BtuJ1 bound to the ligands CNCbl (B_12_) and Cbi. (A) Cartoon representation of BtuJ1 (in rainbow colour, N terminus in blue) bound to Cbi (magenta). (B) Close-up of the residues forming hydrogen bonds (black dashed lines) with Cbi. (C,D) Figure showing a stick representation of CNCbl (B_12_) (C) and Cbi (D).The two axial positions of the cobalt atom are occupied by cyanide (–CN) ligands) (D) bound to BtuJ1 with the 2Fo-Fc electron density contoured at 1.5 σ.

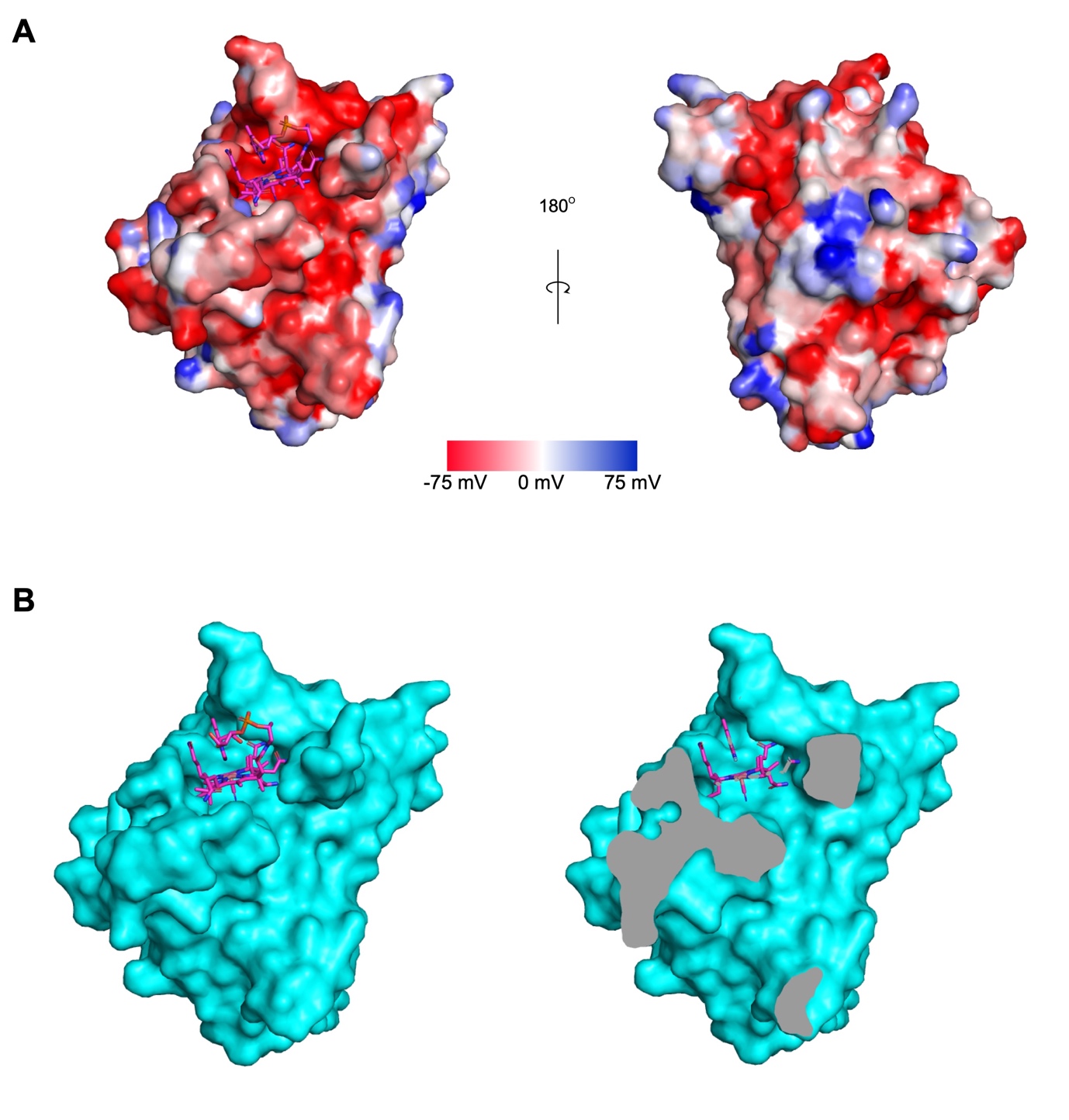

**FIG S2** Analysis of physical properties of the cobamide binding pocket of BtuJ1 (A) Electrostatic surfaces of BtuJ1 generated using Protein-sol (1) B_12_ is represented in magenta. (B) Surface representation of BtuJ1 with CNCbl (B_12_) in magenta (left) and a slice of a surface representation for CNCbl-BtuJ1 showing the cavity for the upper ligand (right).

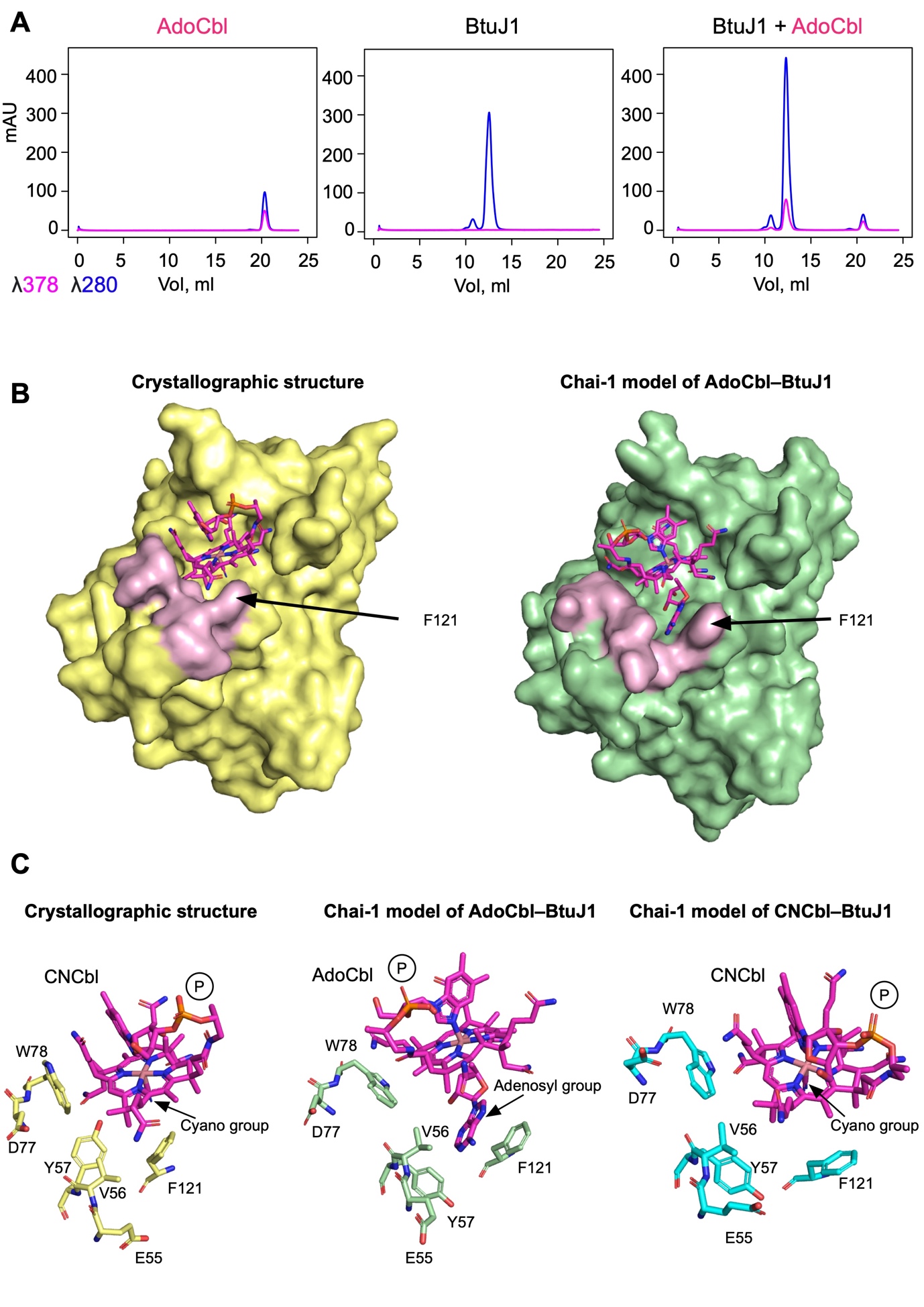

**FIG S3** BtuJ1 binds Adenosylcobalamin (AdoCbl) (A). Size exclusion chromatography (SEC) traces of 5 nanomoles of AdoCbl (top), apo-BtuJ1 (middle) and BtuJ1 incubated with 10 nanomoles of AdoCbl for 1 hour at 20 °C (bottom). Protein absorbance is measured at 280 nm; AdoCbl absorbance is measured at 378 nm. (B) Structural panels showing surface representations of BtuJ1. Left: Crystal structure (yellow) bound to CNCbl (magenta sticks). Right: Chai-1-modelled structure (2) (green) bound to AdoCbl (magenta sticks), in light-pink colour residues that are displaced to enlarge the binding pocket. (C) Comparison of stick representations showing residues displaced to accommodate the adenosyl group. Crystal structure bound to CNCbl (yellow), modelled AdoCbl structure (green) and modelled CNCbl structure (cyan). The phosphate group (P) is shown for all three structures to highlight its displacement. Although the modelled CNCbl structure was expected to match the crystal, there are substantial deviations both in the ligand and protein. For example, the position of Y57 instead aligns more closely with the AdoCbl model, indicating a notable deviation from the experimental structure and underscoring that a need for experimental structure determination remains.

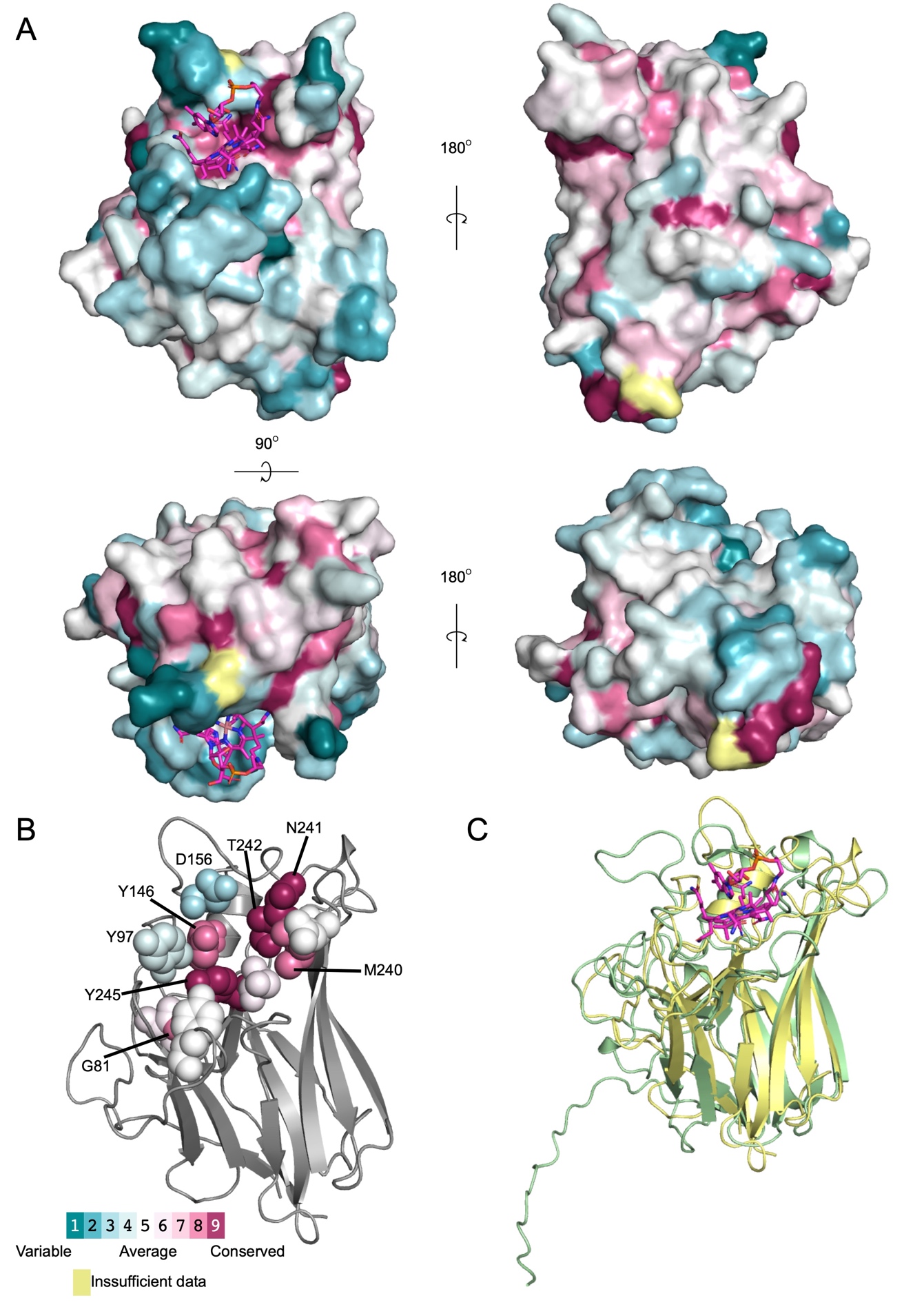

**FIG S4** Sequence and structure conservation of BtuJ1 (A) Views of the surface representation of BtuJ1 coloured with the conservation scores obtained in the ConSurf (3)  analyses. (B) Cartoon representation of BtuJ1 showing the level of conservation of residues implicated in hydrogen bonding (ConSurf). (C) Superposition of the structure of BtuJ1 (yellow) bound to CNCbl (B_12_) and the AlphaFold (4, 5) model for BtuJ2 (green), demonstrating the structural similarity.

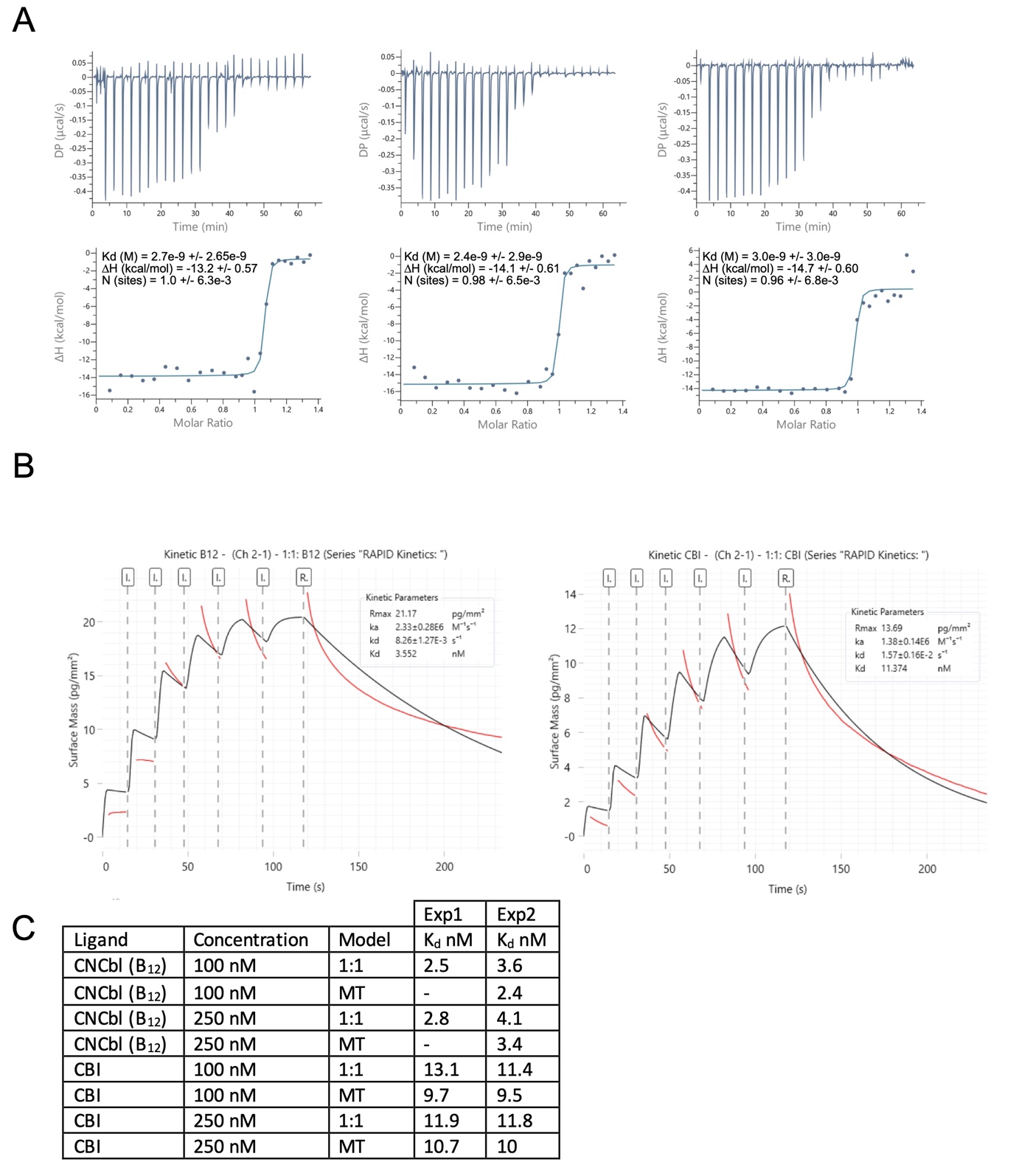

**Fig S5** Ligand binding analysis for BtuJ1 (A) B12 binding to BtuJ1 analysed by ITC. The upper parts of each titration show the ray heats of injection and the lower parts the integrated heats. Data calculated is included into each replica. (B) Grating-Coupled Interferometry (GCI) analysis of BtuJ1 binding to cobinamide (Cbi) and cyanocobalamin (CNCbl). Sensorgrams show surface mass (response units, RU) versus time using the waveRAPID kinetic injection protocol. BtuJ1 was immobilized on a Ni²⁺-NTA sensor surface via His-guided amine coupling. Analytes were injected at 100 or 250 nM. The left panel shows binding to CNCbl; the right panel shows binding to Cbi. Global fitting of the data using a 1:1 binding model is shown in black. Calculated kinetic parameters (association rate k_a_, dissociation rate k_d_, and dissociation constant K_D_) are reported within each panel. Comparable results were obtained via fitting using a mass transport-corrected model (MT), indicating minimal mass transport effects. (C) Table showing the dissociation constants (n = 2) for the analyte at different concentrations and using different analysis models (1:1 and MT).

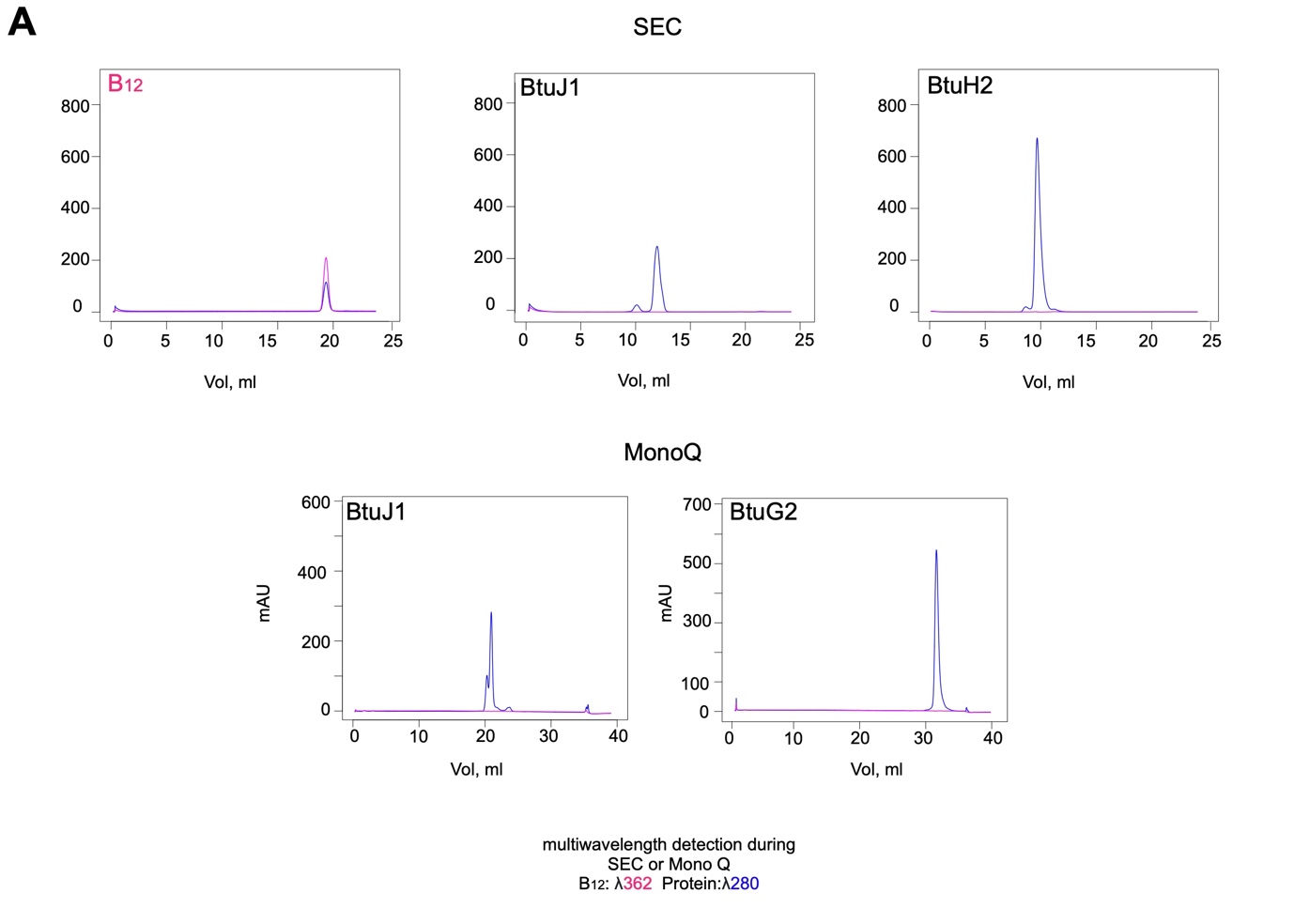
**FIG S6** Controls for the transfer experiment. SEC (top) and MonoQ (bottom) traces of 5 nanomoles of protein or CNCbl (B_12_) incubated 1 hour at 20 °C. Protein absorbance is measured at 280 nm and vitamin B12 absorbance is measured at 362 nm. Data are representative of two independent experiments.

**MOVIE S1**. Electron density map of CNCbl bound to BtuJ1. BtuJ1 is shown in a cartoon representation coloured in a rainbow spectrum, and water molecules are depicted as red spheres.
